# ERK signaling dissolves ERF Repression Condensates in Living Embryos

**DOI:** 10.1101/2021.10.01.462829

**Authors:** Claire J. Weaver, Aleena L. Patel, Stanislav Y. Shvartsman, Michael S. Levine, Nicholas Treen

## Abstract

Phase separation underlies the organization of the nucleus, including the biogenesis of nucleoli and the packaging of heterochromatin. Here we explore the regulation of transcription factor condensates involved in gene repression by ERK signaling in gastrulating embryos of a simple proto-vertebrate (*Ciona*). ERK signaling induces nuclear export of the transcriptional repressor ERF, which has been linked to various human developmental disorders. Using high resolution imaging we show that ERF is localized within discrete nuclear condensates that dissolve upon ERK activation. Interestingly, we observe dynamic pulses of assembly and dissociation during the cell cycle, providing the first visualization of a nuclear phase separation process that is regulated by cell signaling. We discuss the implications of these observations for producing sharp on/off switches in gene activity and suppressing noise in cell-cell signaling events.

## INTRODUCTION

Liquid-liquid phase separation (LLPS) is emerging as a major mechanism for cellular organization (1), including the biogenesis of nucleoli and other components of the nucleus (2, 3). Within nuclei, LLPS has also been implicated in the assembly of the RNA Polymerase II pre-initiation complex at poised promoters (4-6), as well transcriptional silencing. HP1 chromocenters display liquid behavior (7, 8), and the transcriptional repressor Hes.a was recently shown to possess liquid properties when associated with the Groucho (Gro/TLE) corepressor (9). Hes.a/Gro condensates are stable throughout the cell cycle when expressed in living *Ciona* embryos, finally dissolving at the onset of mitosis (9). These observations point towards a simple mechanism for transcriptional repression, whereby Hes.a/Gro condensates exclude transcriptional activators at silent loci (9).

Transcriptional repressors are often deployed to silence gene expression in the absence of cell signaling (10). This default repression is inhibited upon cell signaling to activate gene expression (10, 11). Here, we explore the possibility that cell signaling derepresses gene silencing by influencing the assembly and disassembly of repression condensates. In ERK signaling, the activation of a receptor by its ligand (e.g., FGF) triggers a phosphorylation cascade culminating in the activation and nuclear translocation of ERK (12). Previous studies suggest ERK signaling anagonizes transcriptional repressors such as Capicua, Yan and ERF (13-16), in addition to inducing the activities of transcriptional activators such as ETS and Pointed (17).

In the *Ciona* embryo, FGF/ERK signaling is pervasively used to specify a variety of cell types, including cardiomyocytes and neuronal cell types in the CNS (18-20). This signaling acts upon ETS-class transcription factors - typically interpreted as activators, although de-repression mechanisms are likely to participate as well. The maternally expressed *Ciona* orthologue of the human Ets-2 repressive factor (ERF) is a strong candidate for one such repressor as it persists throughout early development (21). Mammalian ERF is a sequence-specific repressor that recognizes canonical ETS binding sites (15, 22, 23) and is exported from the nucleus upon ERK activation (15, 23). It has been implicated in a variety of human diseases, including craniofacial disorders (24).

We present evidence that ERF forms spherical condensates during *Ciona* embryonic development, but FGF/ERK signaling leads to their dissolution and export from the nucleus. There is a tight correlation between the formation of ERF condensates, association with the Groucho corepressor, and transcriptional silencing of FGF/ERK target genes. Unexpectedly, ERF condensates exhibit a pulse of dissolution and reformation during interphase, suggesting transient rather than sustained FGF/ERK signaling. We suggest that regulating the assembly and disassembly of ERF condensates suppresses noise and produces sharp, switch-like induction of gene expression.

## RESULTS

A survey of the *Ciona* genome identified two putative orthologues of the human ERF repressor. One is maternally expressed in the *Ciona* embryo (Erf.b/KY.Chr4.757), while the other does not produce detectable transcripts until late larval stages (Erf.a/KY.Chr4.415). We focused our efforts on the maternal orthologue, hereafter called ERF.

*Ciona* ERF was fused to mNeongreen (mNg) and expressed in the embryonic ectoderm of gastrulating *Ciona* embryos using a *Sox1/2/3* regulatory sequence (Fig. 1A). Nuclear, punctate fluorescence is readily detected. Super-resolution confocal microscopy revealed punctate distributions of ERF evocative of Hes.a repression condensates (Fig. 1D, Fig. S1A). The corepressor for ERF has not been definitively identified, although there is evidence for potential interactions between human ERF and TLE5 (25). We observed colocalization of Gro (TLE) within ERF puncta (Fig. 1D), suggesting that Gro functions as a corepressor of ERF in the *Ciona* embryo.

**Fig. 1.**
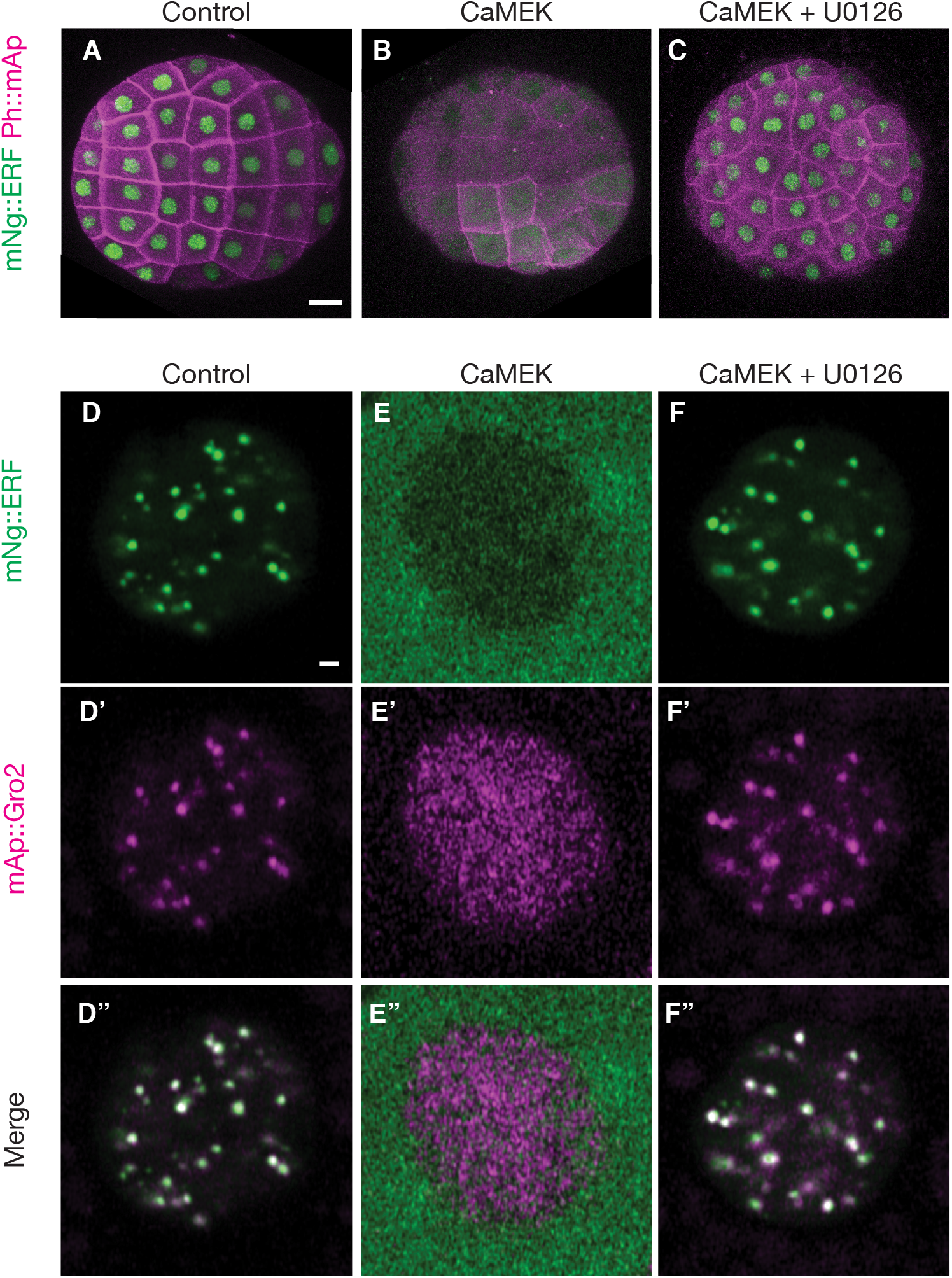
ERF Nuclear export is dependent on ERK phosphorylation. All Figs show ectodermal cells of 110-cell stage expressing electroporated transgenes of fluorescent fusion proteins in ectodermal cells. (A-C) Whole embryo views of live *Ciona* embryos with the indicated experimental treatments. Cell outlines are visualized using PH::mAp. (A) mNg::ERF is localized to the nucleus. Levels of nuclear fluorescence was variable between nuclei. (B) CaMEK overexpression causes ERF to localize outside the nucleus. (C) Treatment of CaMEK overexpressed embryos with the MEK inhibitor U0126 reverses the action of CaMEK and nuclear localization of ERF is restored. (D-F) Confocal sections of single *Ciona* nuclei. mNg::ERF is localized to spherical puncta that colocalize with mAp::Gro2. Ng::ERF is exported to from the nucleus, but mAp::Gro2 remains nuclear when CaMEK is overexpressed. This action is reversed when treated with the MEK inhibitor U0126. Scale bars A: 20 μm, D: 1 μm.

Unlike Hes.a condensates, ERF was erratically distributed across different nuclei, which may be due to localized FGF signaling during embryonic development. To explore this possibility ERK was activated throughout the ectoderm using a variant of human MEK (CaMEK) that possesses greater constitutive activity than commonly used phosphomimetic variants (26). This experimental activation of MEK caused ERF to be excluded from all nuclei (Fig. 1B), demonstrating ERF localization is dependent on ERK signaling. When treated with the MEK inhibitor U0126 ERF nuclear exclusion is reversed and nuclear accumulation is restored (Fig. 1C).

Co-expression of CaMEK not only led to export of ERF from the nucleus, but also caused a loss of ERF/Gro puncta (Fig. 1E). These puncta are restored upon addition of the U0126 inhibitor (Fig. 1F), suggesting that ERF/Gro complexes are specific targets of ERK signaling. By contrast, activation or inhibition of ERK had no noticeable effect on Hes.a condensates (Fig. S1).

Human ERF has an N-terminal ETS DNA binding domain followed by a disordered domain. A region of this disordered sequence (residues 472-530) is essential for repression (Fig. 2A, 15). The *Ciona* homologue of ERF has a similar N-terminal ETS domain, but also contains an additional 139 amino acid (AA) residues in the C-terminus (Fig. 2B; see below). This region is less disordered than its human counterpart (Fig. 2B). It was previously shown that mutations in the DNA binding of Hes.a resulted in condensates exhibiting definitive liquid-like properties (9). We similarly observe that a K83Q mutation in the ERF ETS domain forms fewer, larger puncta, that retain association with Gro (Fig. 2C). This mutation mimics a human clinical variant associated with craniosynostosis (24). The mutant condensates display similar responses to activated or inhibited MEK as their wildtype counterparts (Fig. 2D, E), suggesting that responses to ERK signaling can occur irrespective of DNA binding. They also exhibit fusions of individual puncta, consistent with the formation of ERF/Gro repression condensates via LLPS (Fig. 2F, Supplemental Movie 1).

**Fig. 2.**
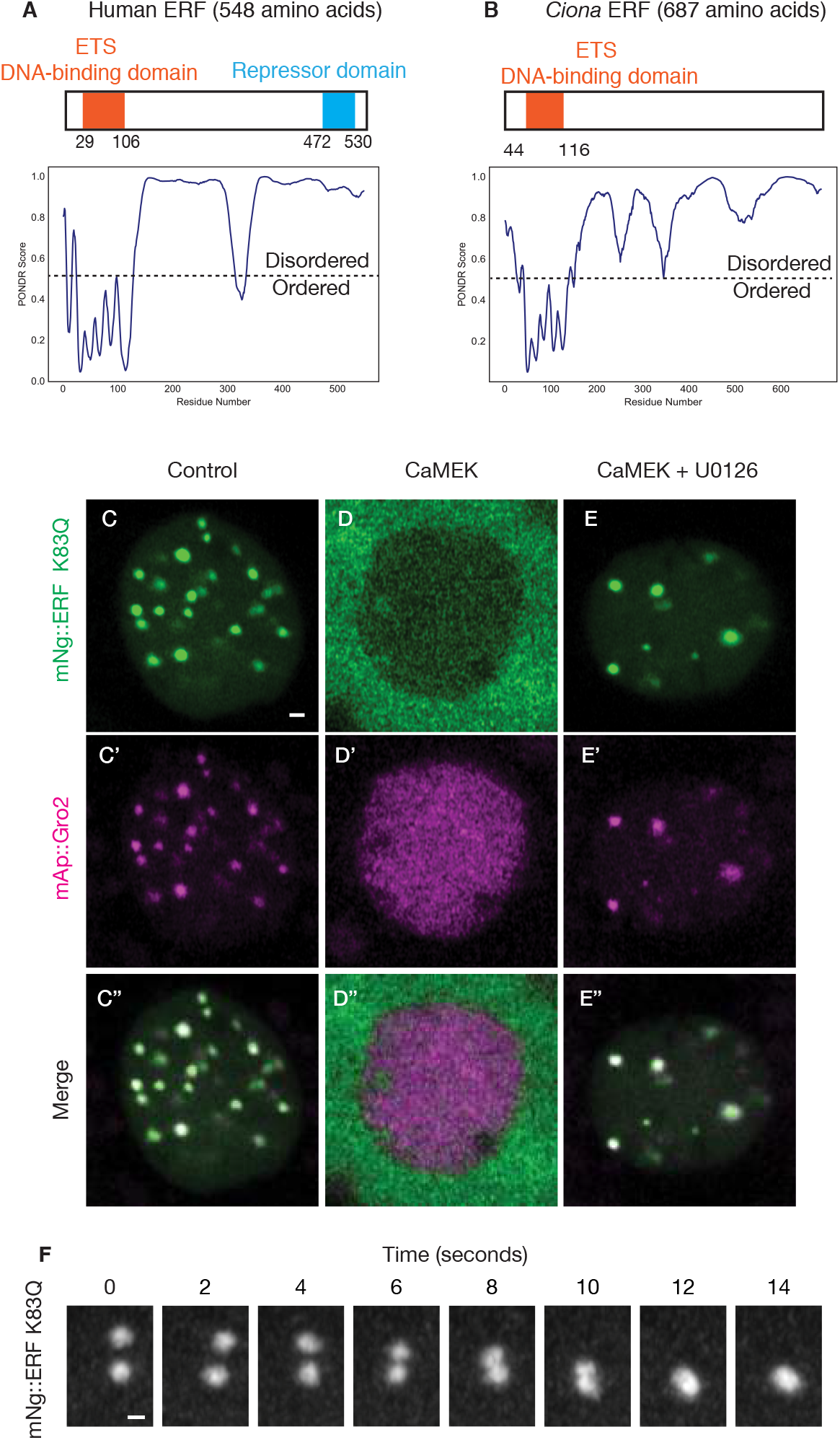
ERF DNA-binding mutants show liquid properties. (A,B) Primary structures of Human and Ciona ERF showing the ETS DNA-binding domain in orange and the repressor domain in blue. Below the structures are PONDR plots showing the predicted ordered and disordered regions. (C-E) Confocal sections of single *Ciona* nuclei. mNg::ERF K83Q is localized to spherical puncta that colocalize with mAp::Gro2. Ng::ERF K83Q is exported to from the nucleus, but mAp::Gro2 remains nuclear when CaMEK is overexpressed. This action is reversed when treated with the MEK inhibitor U0126. (F) Maximum intensity confocal projection showing the fusion of 2 Ng::ERF K83Q droplets over 14 seconds. Scale bars C: 10 μm, D: 0.5 μm.

To identify a putative repression domain, we created a series of truncations of ERF. Removal of the C-terminal 140 amino acid residues disrupted droplet formation, while loss of an additional 20 AA residues resulted in uniform distribution throughout the nucleus (Fig. S2A). This behavior is similar to previous observations for Hes.a where the removal of the Gro-interaction motif (WRPW) abolished droplet formation (9). Based on these observations we designate the C-terminal 160 residues of Ci-ERF as a droplet forming domain (Fig. S2B).

We next asked whether this domain mediates repression. Chimeric proteins that replace the DNA binding domain of human ERF with the DNA binding domains of other transcription factors were found to retain ERK-modulated repression activities (15). We therefore replaced the ETS DNA binding domain of *Ciona* ERF with the bHLH DNA binding domain of Hes.a (Fig. 3A). The resulting chimera forms nuclear puncta (Fig. S3A), displays colocalization with Gro, and is imported and exported from nuclei in response to experimental manipulation of MEK (Fig. S3B,C). Removal of the 160 AA droplet domain from this chimeric protein resulted in a loss of droplets and Gro colocalization (Fig. 3B-D).

**Fig. 3.**
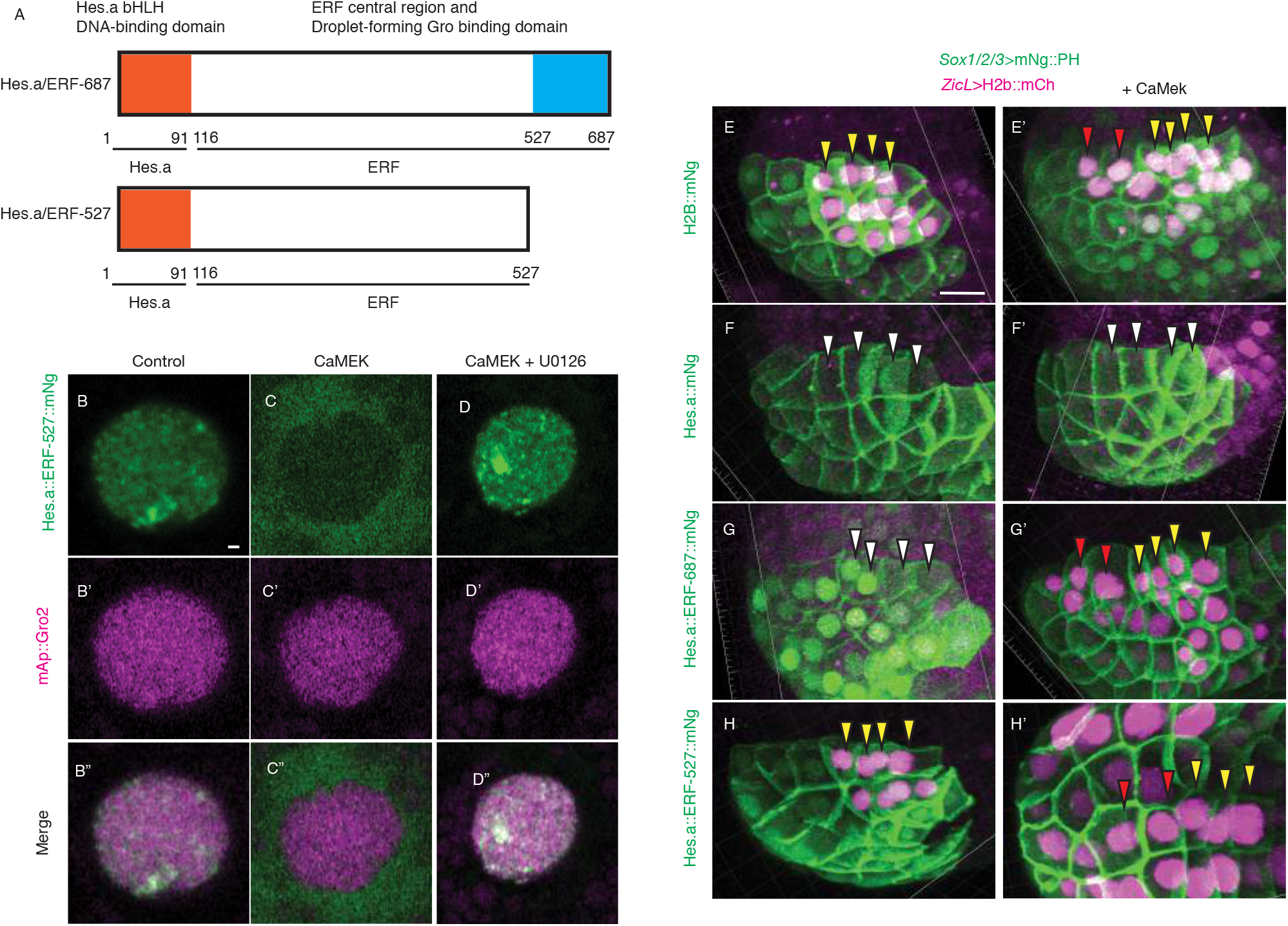
*Ciona* ERF contains a C-terminal repressor domain. (A) Primary structures of Hes.a/ERF chimera proteins with the DNA binding and droplet froming domains indicated. (B-D) Confocal sections of single *Ciona* nuclei. Hes.a::ERF-527::mNg is localized to puncta that do not colocalize with mAp::Gro2. Hes.a::ERF-527:mNG is exported to from the nucleus, but mAp::Gro2 remains nuclear when CaMEK is overexpressed. This action is reversed when treated with the MEK inhibitor U0126. (E,E’) Expression of ZicL reporter in the *Ciona* neural plate is expanded anteriorly when CaMEK is expressed. (F,F’) Hes.a::mNG represses all ZicL expression in the neural plate. (G,G’) Hes.a::ERF-687::mNG represses ZicL. This repression is inactivated by CaMEK. (H,H’) Hes.a::ERF-527::mNg is unable to repress ZicL. Yellow arrowheads indicate Wild type expression. Red arrowheads indicate ectopic expression, white arrowheads indicate repression of wild type expression. Cell outlines are indicated with Ph::mNG as in several cases the transcription factors of interest are degraded by the time reporter activity is detectable. Scale bars A: 1 μm, E: 015 μm.

The Hes.a/ERF chimera appears to repress a Hes.a target gene in response to FGF/ERK signaling. The Hes.a target gene ZicL (27) is expressed in neuronal precursor cells of rows I-IV of the presumptive neural plate (28). A reporter containing the upstream regulatory sequences of ZicL recapitulates this localized pattern of expression (Fig. 3D, 20). Co-expression of CaMEK throughout the early ectoderm results in an anterior expansion of the ZicL reporter gene (Fig. 3E’). Hes.a has previously been shown to repress this reporter (Fig. 3F, 9), and it continues to do so even upon coexpression of CaMEK (Fig. 3F’). By contrast, the Hes.a/Erf chimera represses ZicL in the neural plate, but not in posterior cells that are adjacent to a localized source of FGF/ERK signaling (Fig. 3G, 29). Moreover, repression is completely abolished when CaMEK is expressed throughout the neural plate (Fig.4G’). These observations suggest that the C-terminal 160 residues of the ERF protein are required for repression by recruiting Gro and derepression by FGF/ERK signaling (Fig. 3H).

Live imaging of gastrulating embryos revealed dynamic import and export of ERF from the nucleus over the course of a single interphase (Fig. 4A, S4, Supplemental Movies 2, 3). This behavior was variable, with some cells having ERF remain nuclear while others displayed complete exclusion from the nucleus throughout interphase (Supplemental Movies 2,3). ERF condensates were limited to ∼500nm in diameter even upon complete inhibition of ERK signaling with U0126 (Fig. 4, Supplemental Supplemental Movies 4,5). Consistent with previously observed Hes.a condensate behaviors (9), the DNA-binding mutant formed fewer and larger droplets (Fig.S4, Supplemental Movies 4,). Pulsatile behavior was still observed, although the initial formation of condensates was short lived (Fig. S4, Supplemental Movies 6,7). When treated with U0126, mutant ERF condensates continued to grow and fuse to sizes over 1uM, considerably larger than wild-type condensates (Figs 4, S4, Supplemental Movies 8,9). These observations suggest that ERF condensate growth is limited by both DNA binding and ERK phosphorylation. This pulsatile behavior was never observed when embryos were treated with the U0126 inhibitor (Fig.4D), suggesting that nuclear export and dissolution is due to ERK signaling.

**Fig. 4.**
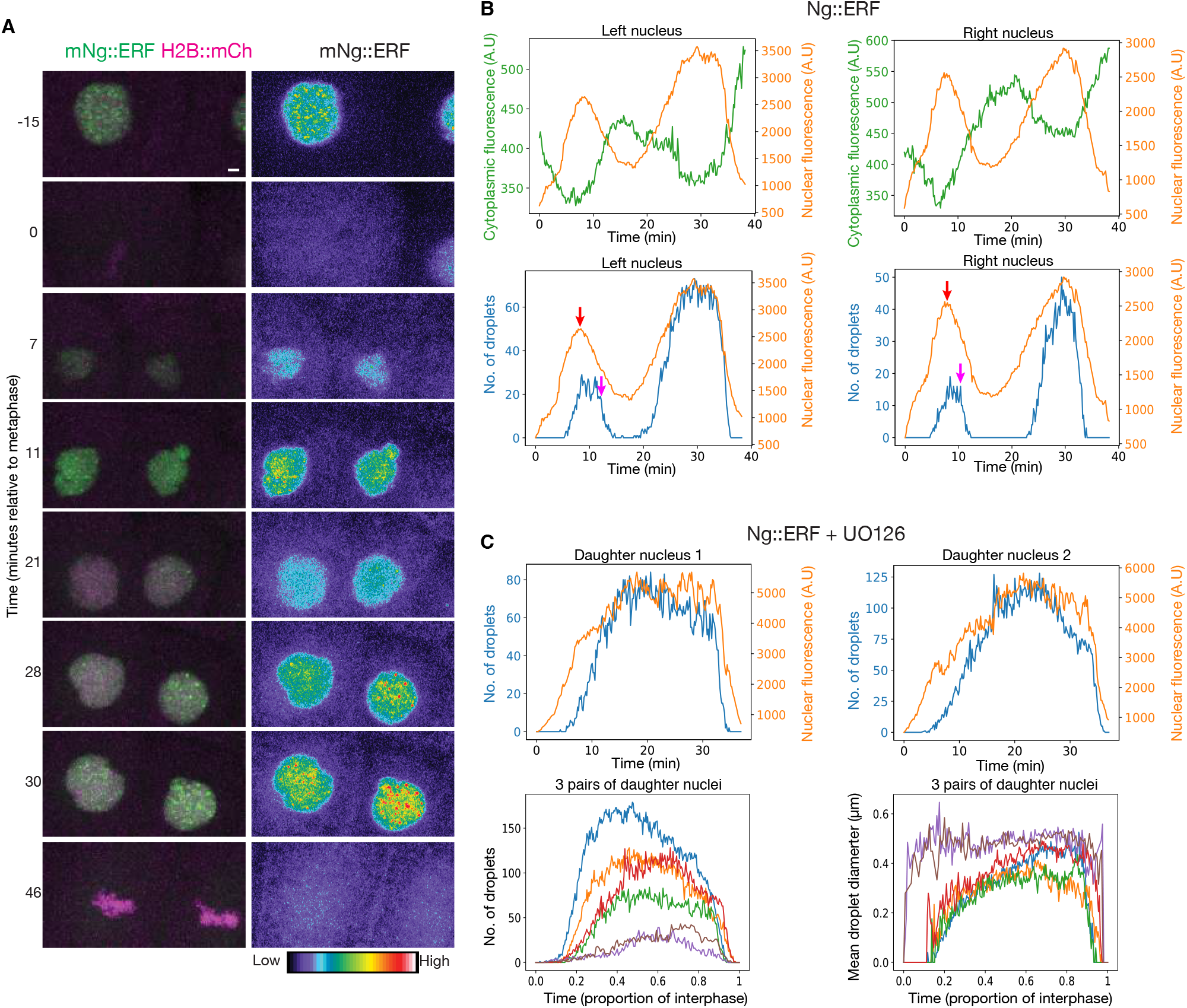
ERF pulses during interphase. (A,B) Maximum intensity confocal projection of a *Ciona* nucleus dividing and the daughter cells throughout interphase. Ng::ERF is exported and imported from the nucleus throughout interphase. (C) Quantifications of the nuclear fluorescence intensity, number of droplets, and cytoplasmic fluorescence intensity of the daughter nuclei in A and B. Red arrowheads indicate the approximate time that nuclear fluorescence intensity decreases and yellow arrowheads indicate the approximate time that number of droplets abruptly decreases. Scale bar: 1 μm.

We observed that nuclear export is accompanied by a decrease in nuclear fluorescence and a concurrent increase in cytoplasmic fluorescence (Fig. 4 C). While nuclear fluorescence decreases, droplet number remains relatively unchanged for 3-5 mins. After this lag, there is a sudden and uniform dissolution of ERF/Gro condensates (Fig.4C, S3), suggesting ERK-mediated export precedes dissolution. This behavior is not observed at mitosis, where the decrease in nuclear florescence and dissolution of droplets were tightly correlated (Fig.4, S3). We discuss the implications of these observations below.

## DISCUSSION

We have presented evidence that ERF associates with Gro and forms repression condensates. This association depends on an extended C-terminal region that is required for the recruitment of Gro, the formation of condensates, and transcriptional repression. This differs from the short, dedicated WRPW motif that mediates Hes.a-Gro associations, raising the possibility of distinct modes of condensate formation for Hes.a and ERF. A particularly striking finding of our analysis is the pulsatile assembly and dissolution of ERF/Gro condensates during the cell cycle. These dynamics sharply contrast with the assembly of stable Hes.a condensates, highlighting the role of FGF/ERK signaling in the precision of gene activity.

ERK signaling pathways often display oscillatory or pulsatile dynamics (30, 31). However, this behavior was not anticipated for *Ciona* embryos since all signaling events, including FGF signaling, is characterized by stable direct cell-cell contacts (18, 32, 33). Nonetheless, we observe clear pulses of ERF/Gro assembly and disassembly during the cell cycle. These observations suggest a short permissive period for response to ERK signaling during the cell cycle, despite sustained cell-cell contacts. This restrictive period of response might suppress noise and help ensure switch-like induction of gene activity by extracellular signals such as FGF.

Previous studies have implicated Ephrin is a key antagonist of FGF signaling in *Ciona* (34,35), raising the possibility that Ephrin inhibition works in concert with LLPS to ensure transcriptional precision. We note, however, that Ephrin activities are mainly detected at different stages of development than those examined in this study. Future studies will explore the relative roles of LLPS and Ephrin in the control of gene activity by FGF and other mediators of ERK signaling.

How does ERK signaling trigger the dissociation of ERF/Gro condensates? We tested the possibility that it might be due to ERF nuclear export. The idea is that export would trigger a domino effect, whereby ERF in the condensed phase would be released to compensate for reduced concentrations of ERF in the dilute phase. To test this idea we treated embryos with Leptomycin B, a drug shown to inhibit export of mammalian ERF (23). ERF/Gro condensates appear to dissolve on schedule despite this block in ERF nuclear export (Fig.S6). These observations are consistent with fast de-repression of genes in the *Drosophila* embryo, which occurs before ERK signals lead to nuclear export of the Capicua repressor (13, 36).

We therefore suggest that ERK-mediated phosphorylation underlies the dissolution of ERF repression condensates (Fig.5). It remains to be determined whether this phosphorylation occurs in the dilute phase, the condensed phase, or both. Regardless of mechanism, the dissolution of repression condensates is an inherently effective mechanism for the rapid, switch-like induction of gene expression in response to cell-cell signaling. It seems likely that a similar strategy is used for a variety of developmental patterning processes.

**Fig. 5.**
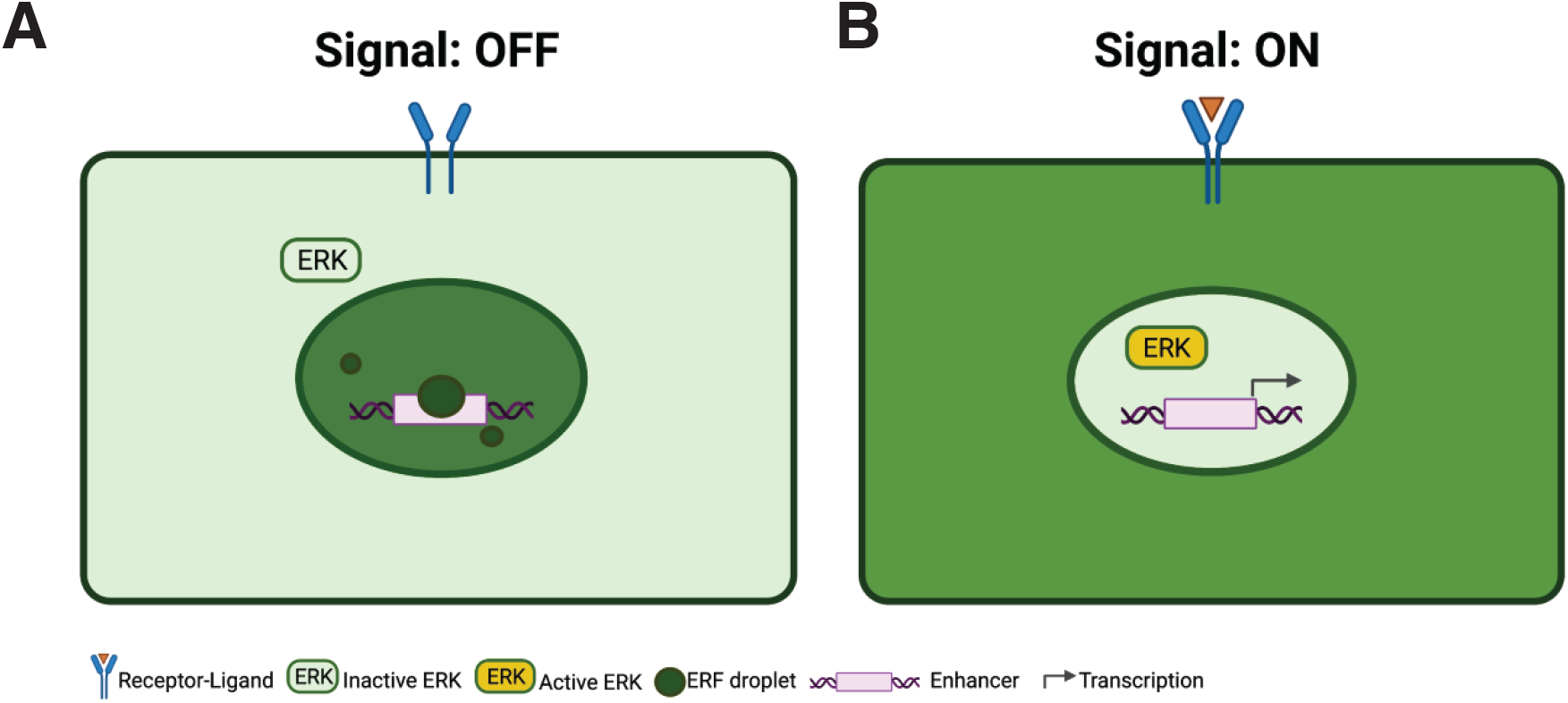
ERF derepression by dissolution of repressive condensates. (A) In the absence of an inductive signal ERK is inactive and ERF is localized to the nucleus where it forms repressive condensates as well as a dilute phase. (B) When a signal is activated, ERK enters the nucleus and will immediately phosphorylate ERF, causing it to be exported from the nucleus and for repressive condensates to dissolve. After an extended or intense period of signaling the repressive condensates will dissolve and transcriptional activation can proceed.

---

**Supplementary Fig. 1.**
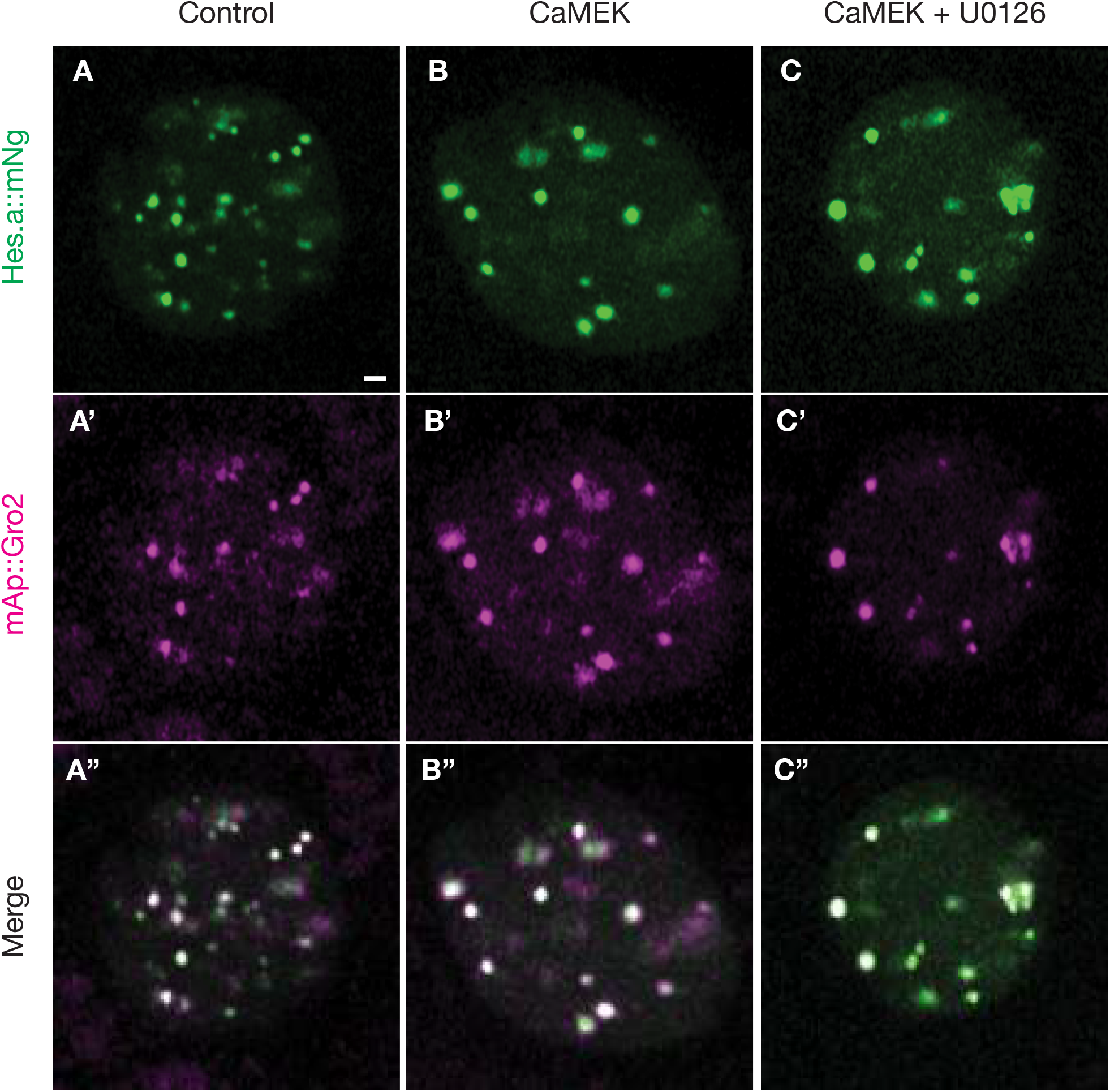
Hes.a does not respond to ERK activity. (A-C) Confocal sections of single *Ciona* nuclei. Hes.a::Ng is distributed in spherical puncta that colocalize with mAp::Gro2 in all conditions. Scale bar: 1 μm.

**Supplementary Fig. 2.**
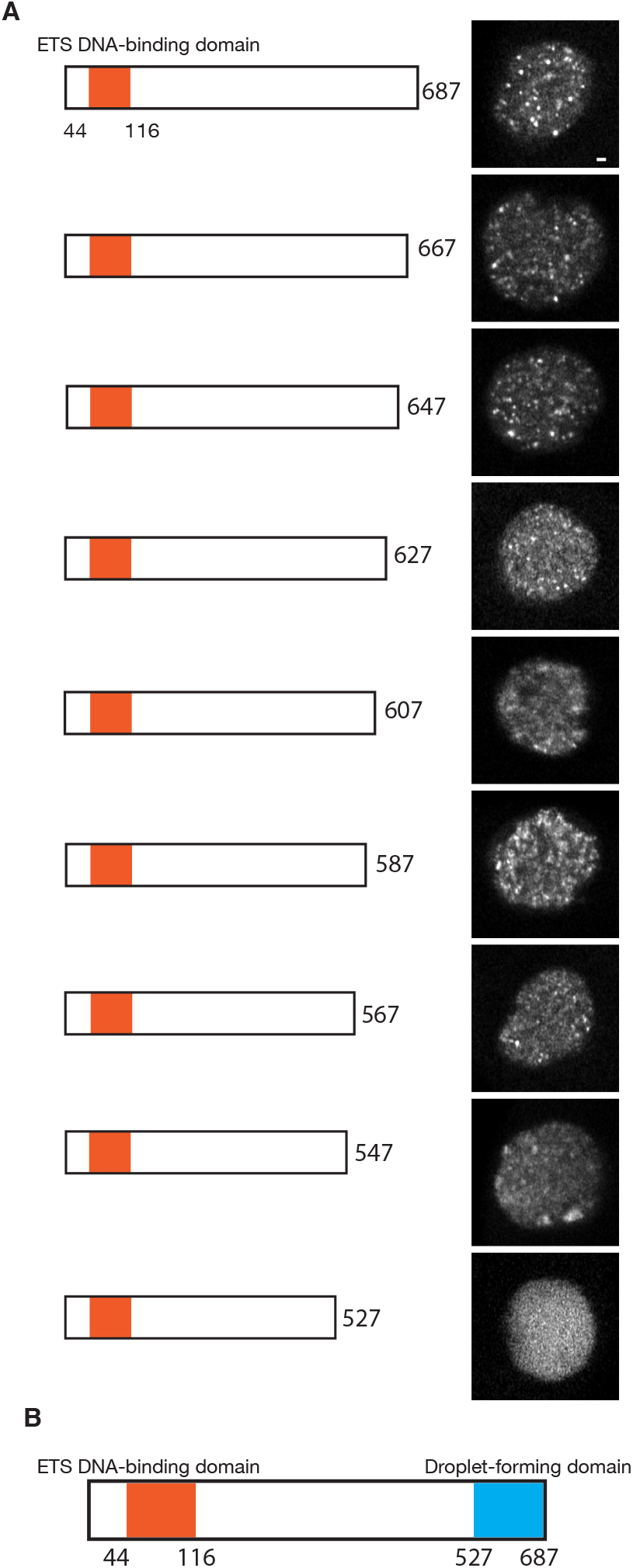
The C-terminus of ERF is a droplet forming domain. (A) Primary structure of *Ciona* truncated ERF showing the ETS DNA-binding domain in orange and indicating the total length of the protein. Corresponding images show confocal sections of single *Ciona* nuclei expressing the truncated form of ERF. Some droplet formation can be seen until 160 C-terminal residues are removed. (B) Primary structure of *Ciona* ERF showing the ETS DNA-binding domain in orange and the droplet forming domain in blue. Scale bar: 1 μm.

**Supplementary Fig. 3.**
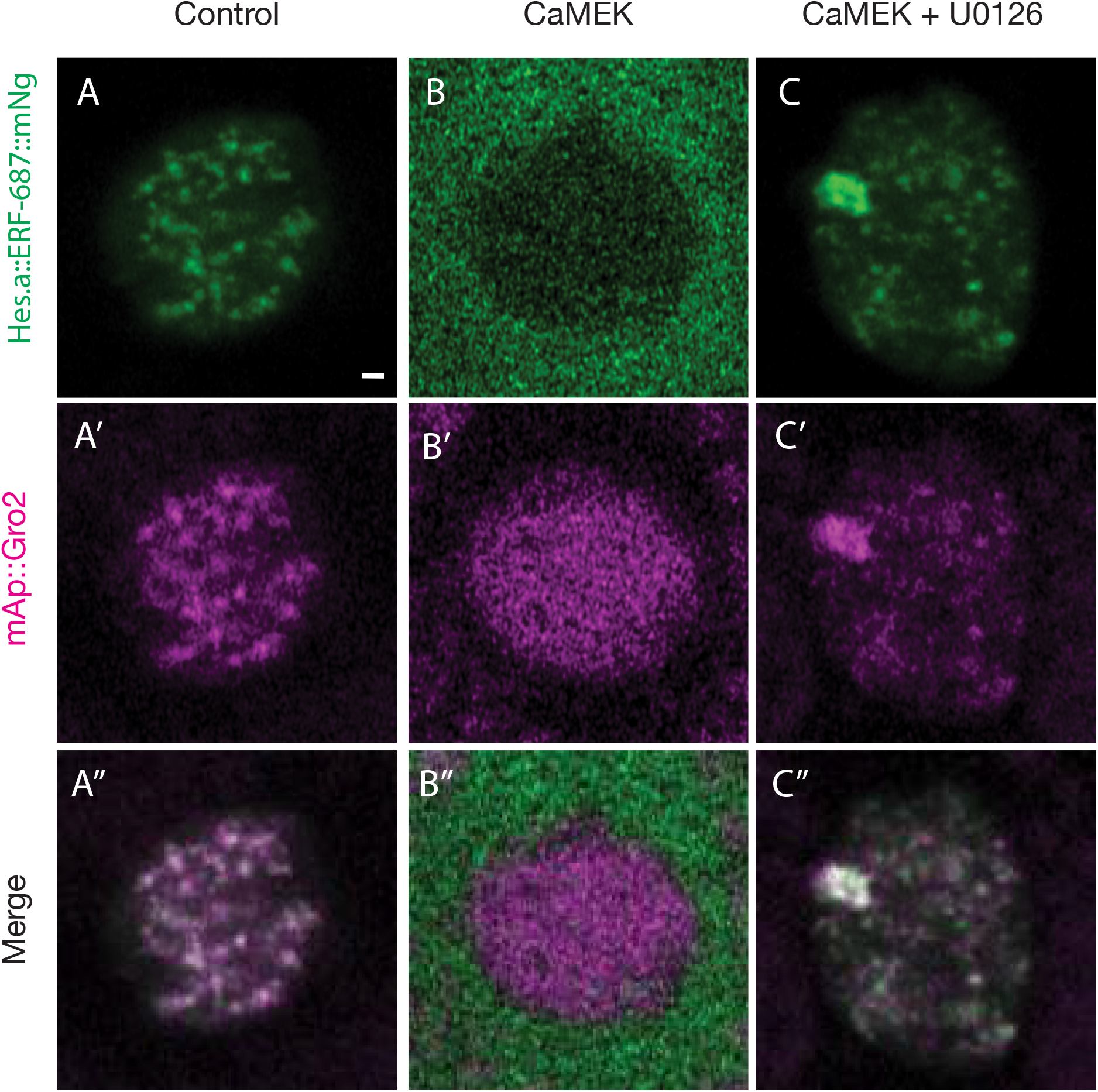
A Hes.a-ERF fusion shows droplet-like distribution and ERK phosphorylation dependent nuclear export. (A-C) Confocal sections of single *Ciona* nuclei. Hes.a::ERF-687::mNg is localized to puncta that colocalize with mAp::Gro2. Hes.a::ERF-687:mNG is exported to from the nucleus, but mAp::Gro2 remains nuclear when CaMEK is overexpressed. Treatment with U0126 restores nuclear localization and puncta formation. Scale bar: 1 μm.

**Supplementary Fig. 4.**
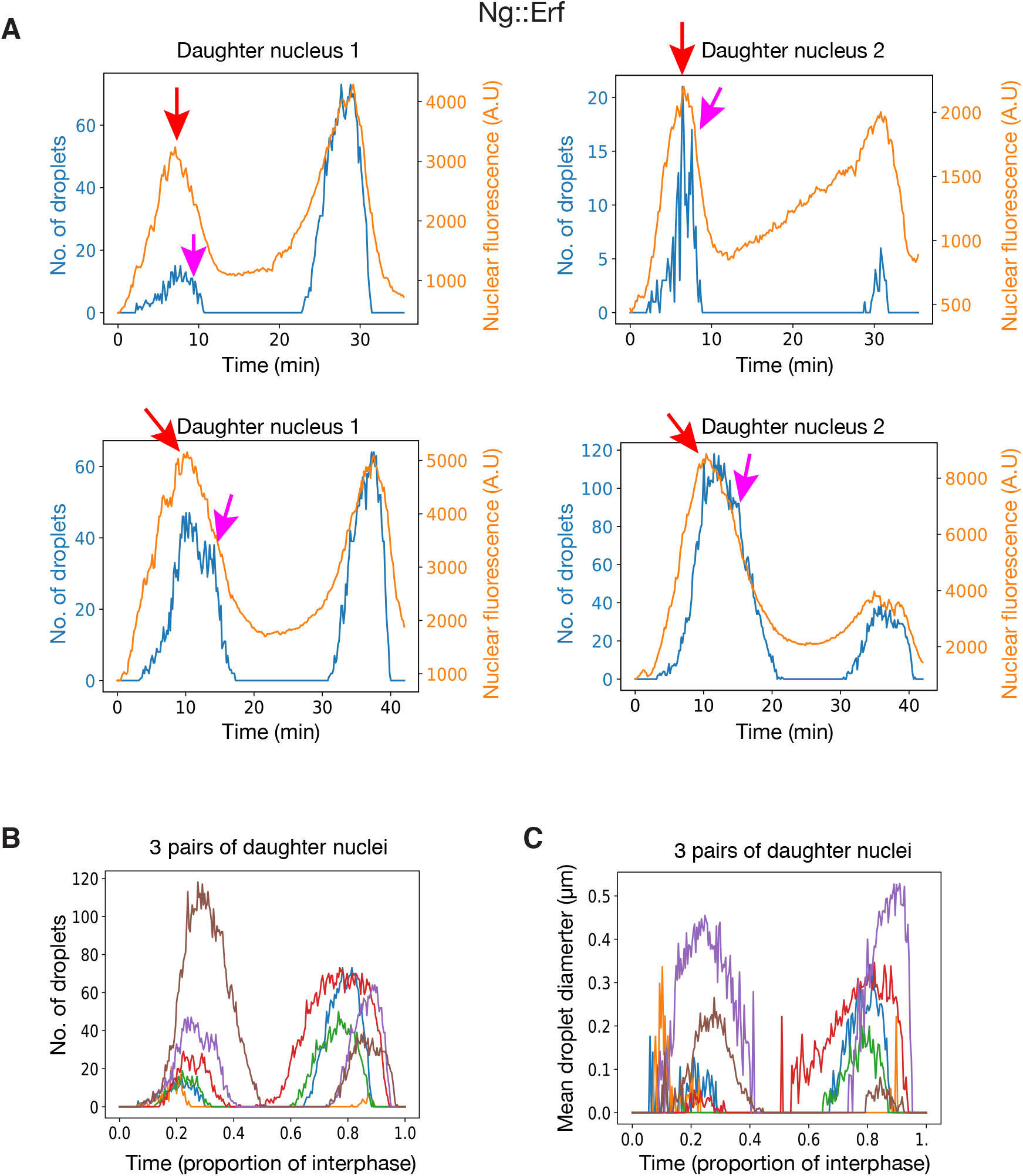
Additional examples of ERF pulses during interphase. (A) Quantifications of the Ng::ERF nuclear fluorescence intensity and number of droplets intensity of 2 pairs of sister nuclei. Red arrows indicate the approximate time that nuclear fluorescence intensity decreases, and magenta arrows indicate the approximate time that number of droplets abruptly decreases. (B) Quantifications of number of Ng::ERF droplets from the 6 nuclei in (A) and Fig. 4B. Time is normalized to the relative proportion of interphase. (C)

**Supplementary Fig. 5.**
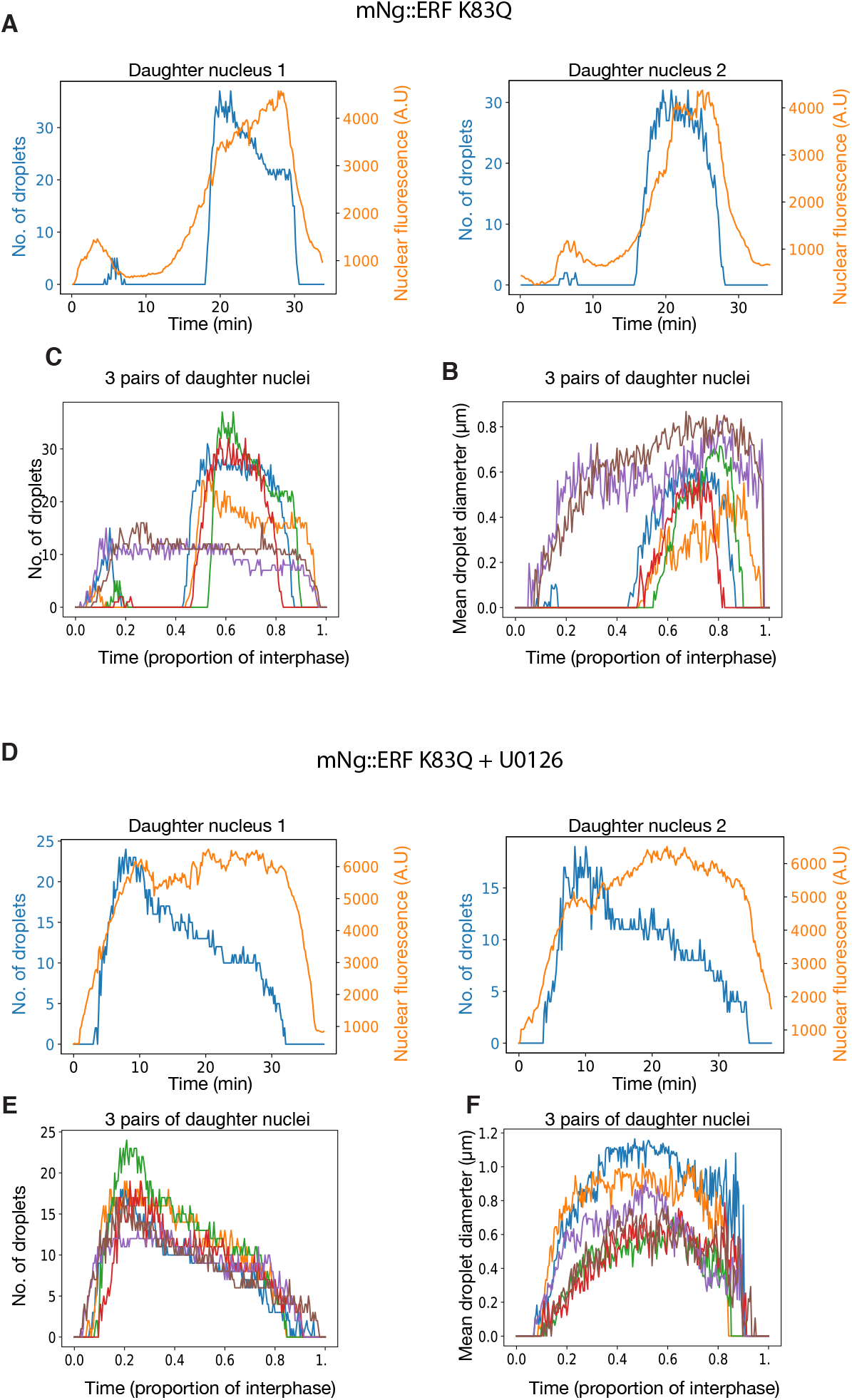
Mutant ERF shows altered droplet behaviors. (A) Quantifications of the Ng::ERF K83Q mutant nuclear fluorescence intensity and number of droplets intensity for a pair of sister nuclei. (B) Quantifications of the number of droplets for the Ng::ERF K83Q mutant in 6 pairs of sister nuclei including the pair shown in A. (C) Quantifications of the mean droplet diameter for the Ng::ERF K83Q mutant in 6 pairs of sister nuclei including the pair shown in A. (D-F) The same conditions as A-C but experiments were performed in the presence of the inhibitor U0126.

**Supplementary Fig. 6.**
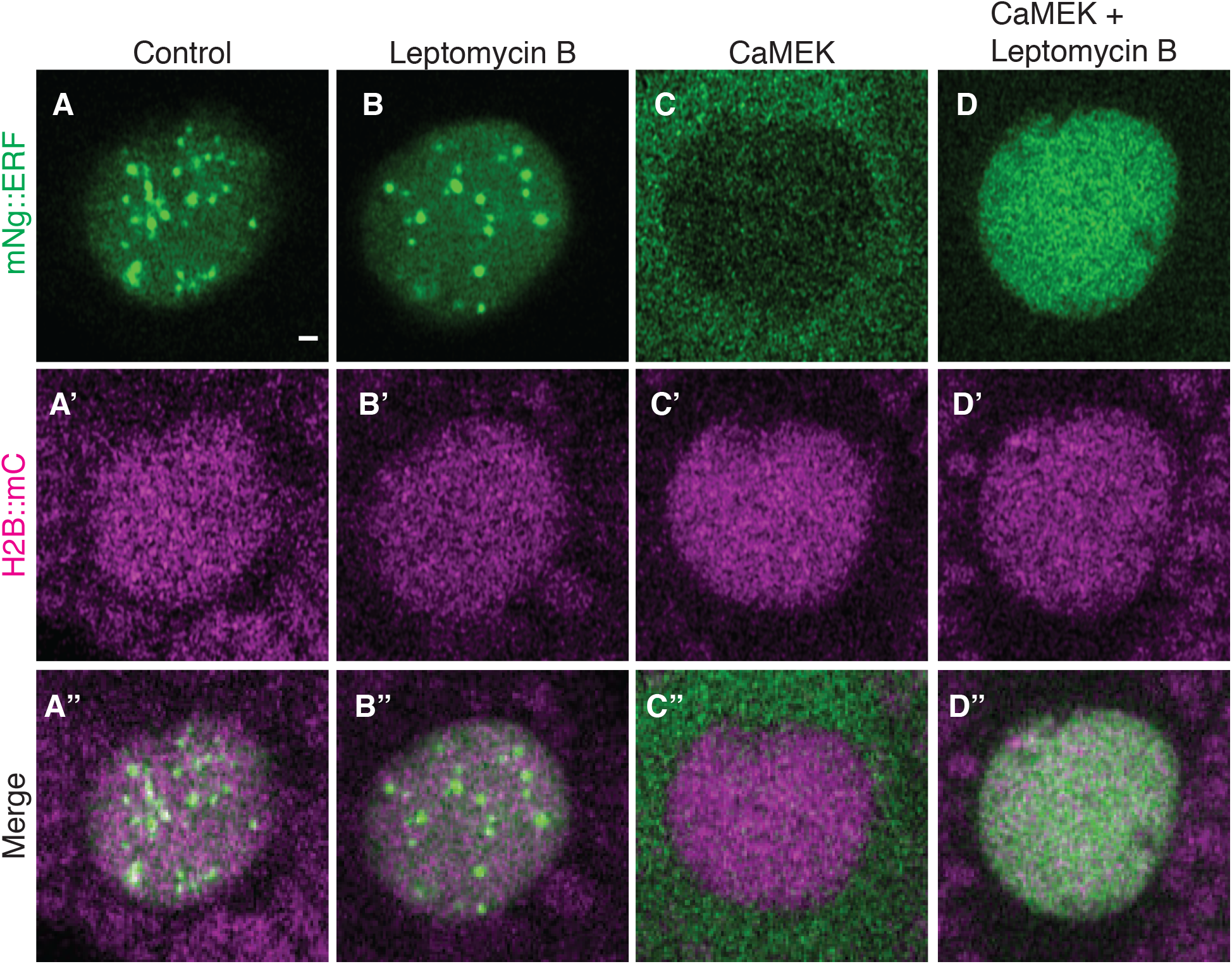
ERF droplets do not require nuclear export to dissolve. (A-D) Confocal sections of single *Ciona* nuclei expressing Ng::ERF and H2B::mC. (A) control embryos show nuclear droplets. (B) ERF Droplets persist when embryos are treated with LeptomycinB. (C) ERF Droplets dissolve and ERF is exported from the nucleus when MEK is activated. (D) Nuclear ERF is recovered, but droplets do not form when MEK is activated and embryos are treated with Leptomycin B. Scale bar: 1 μm.

**Supplementary movie 1. Fusion of mNg::ERF K83Q droplets**.

**Supplementary movie 2. mNG::ERF in living *Ciona* embryos**.

**Supplementary movie 3. 3 separate mitoses depicting mNG::ERF pulses in living *Ciona* embryos**.

**Supplementary movie 4. mNG::ERF in living *Ciona* embryos treated with U0126**.

**Supplementary movie 5. A single mitosis depicting mNG::ERF in living *Ciona* embryos treated with U0126**.

**Supplementary movie 6 mNg::ERF K83Q DNA binding mutant in living *Ciona* embryos**.

**Supplementary movie 7. A single mitosis depicting mNG::ERF K83Q DNA binding mutant in living *Ciona* embryos**.

**Supplementary movie 8 mNg::ERF K83Q DNA binding mutant in living *Ciona* embryos treated with U0126**.

**Supplementary movie 9. A single mitosis depicting mNG::ERF K83Q DNA binding mutant in living *Ciona* embryos treated with U0126**.

## METHODS

### Animals

Wild *Ciona intestinalis* (Pacific populations, Type A, also referred to as *Ciona robusta*) adults were commercially sourced from San Diego County, Ca by M-Rep. Animals were kept in aerated artificial seawater at 18°C. U0126 treatment concentration was 8 μM, Leptomycin B treatment concentration was 0.2 μM. Control embryos were treated with either 0.1% DMSO or ethanol.

### Molecular Cloning

All novel plasmids constructed for this study were based on the *pSP Sox1/2/3>* plasmid previously described (9) with the open reading frames replaced by PCR amplifications using a proofreading DNA polymerase (Primestar, Takara) and plasmids were assembled from linear PCR products using NEBuilder HiFi DNA Assembly Master Mix (New England Biolabs). Full plasmid sequences and descriptions of the individual cloning steps can be provided upon request. The *ZicL>H2B::mCherry* plasmid has previously been described (29). The constitutively active variant of Human MEK has been previously described (26).

### Electroporations

Dechorionated *Ciona* zygotes were electroporated as previously described (9). All experiments were replicated at least in triplicate using different batches of *Ciona* eggs.

### Imaging

Imaging was performed as previously described (9) using a Zeiss 880 Confocal microscope equipped with an Airyscan detector in fast mode. Exact settings and raw imaging files are available upon request.

### Image Analysis

All images were Airyscan processed using Zeiss Zen software (ZEN Version 2.3 and 2.6, Zeiss) The identification and quantification of droplets from confocal images were performed using the *Imaris* spot detection function. All conditions were quantified using the green fluorescence channel across all conditions. Estimated sizes of droplets ranged from 0.2-0.3 um (WT and mutant ERF) and 0.5 um (inhibitor treatments). Background subtraction and region growing were accounted for throughout the cell cycle to accurately report droplet quantity and size. Droplet number in Imaris was filtered using the quality filter and setting a droplet threshold of 95% to prevent false droplet identification. The absolute intensity was used to determine spot regions/diameter. All statistics were exported from Imaris. Further processing was done in Python to fill in the info for frames without droplets as containing 0 droplets of diameter 0. All values exactly equal to the cutoff were excluded from further analysis. The remaining values were normalized to cell cycle length and plotted using matplotlib.

Nuclear and cytoplasmic fluorescence quantification of Ng::ERF was performed in Fiji. A nuclear and cytoplasmic mask was defined and average fluorescence intensities were recorded. Data was plotted alongside droplet data using matplotlib.

## ACKNOWLEDGEMENTS

We thank Clifford Brangwynne, Ned Wingreen and members of the Levine and Brangwynne labs for helpful discussions. We thank Chris Killen for video editing. This research was funded by an NIH grant (NS076542) to M.S.L. N.T is funded by a Princeton Catalysis Initiative grant. C.J.W. is funded by a National Science Foundation graduate research fellowship

## Notes

### Competing Interest Statement

The authors have declared no competing interest.

